# Immune Checkpoint Blockade Delays Cancer and Extends Survival in Murine DNA Polymerase Mutator Syndromes

**DOI:** 10.1101/2024.06.10.597960

**Authors:** Akshada Sawant, Fuqian Shi, Eduardo Cararo Lopes, Zhixian Hu, Somer Abdelfattah, Jennele Baul, Jesse Powers, Christian S. Hinrichs, Joshua D. Rabinowitz, Chang S. Chan, Edmund C. Lattime, Shridar Ganesan, Eileen White

**Affiliations:** Rutgers Cancer Institute of New Jersey, Rutgers University, New Brunswick, NJ 08903, USA; Ludwig Princeton Branch, Ludwig Institute for Cancer Research, Princeton University, Princeton, NJ 08544, USA; Rutgers Robert Wood Johnson Medical School, Piscataway, NJ 08854, USA; Department of Molecular Biology and Biochemistry, Piscataway, NJ 08854, USA

## Abstract

Mutations in polymerases *Pold1* and *Pole* exonuclease domains in humans are associated with increased cancer incidence, elevated tumor mutation burden (TMB) and response to immune checkpoint blockade (ICB). Although ICB is approved for treatment of several cancers, not all tumors with elevated TMB respond. Here we generated *Pold1* and *Pole* proofreading mutator mice and show that ICB treatment of mice with high TMB tumors did not improve survival as only a subset of tumors responded. Similarly, introducing the mutator alleles into mice with Kras/p53 lung cancer did not improve survival, however, passaging mutator tumor cells *in vitro* without immune editing caused rejection in immune-competent hosts, demonstrating the efficiency by which cells with antigenic mutations are eliminated. Finally, ICB treatment of mutator mice earlier, before observable tumors delayed cancer onset, improved survival, and selected for tumors without aneuploidy, suggesting the use of ICB in individuals at high risk for cancer prevention.

**Highlights:**

- Germline somatic and conditional *Pold1* and *Pole* exonuclease domain mutations in mice produce a mutator phenotype.
- Spontaneous cancers arise in mutator mice that have genomic features comparable to human tumors with these mutations.
- ICB treatment of mutator mice with tumors did not improve survival as only a subset of tumors respond.
- Introduction of the mutator alleles into an autochthonous mouse lung cancer model also did not produce immunogenic tumors, whereas passaging mutator tumor cells *in vitro* caused immune rejection indicating efficient selection against antigenic mutations *in vivo*.
- Prophylactic ICB treatment delayed cancer onset, improved survival, and selected for tumors with no aneuploidy.

**Graphical Abstract:** 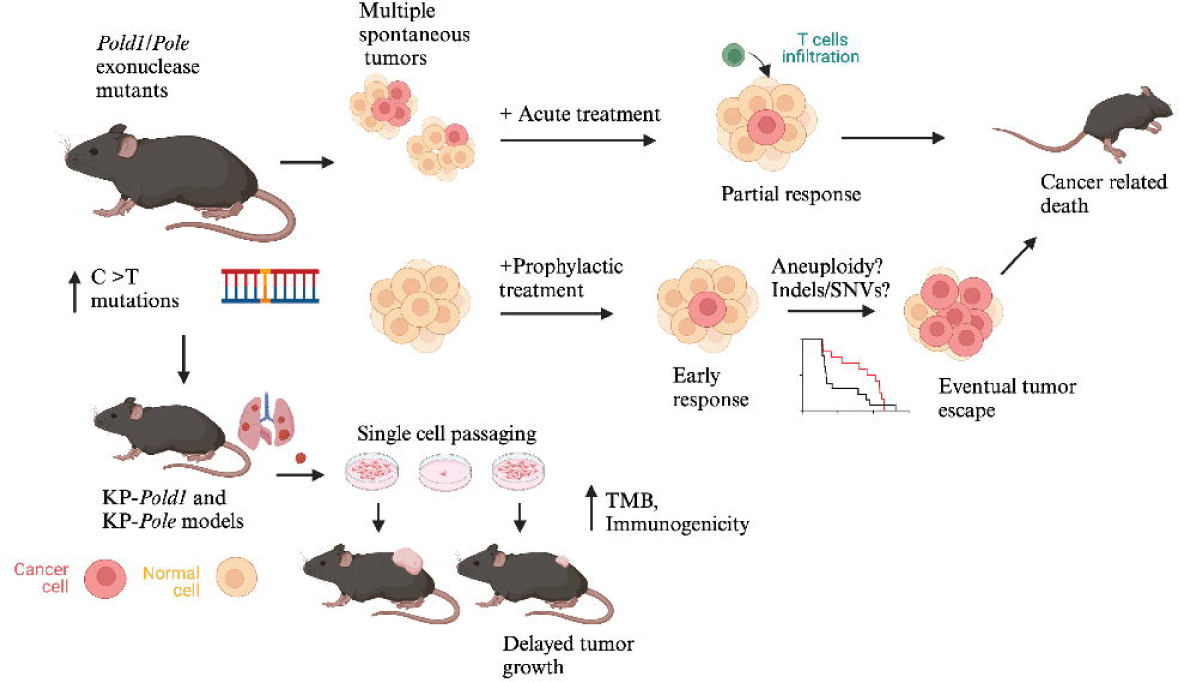

## Introduction

Immune checkpoint blockade (ICB) has provided an extraordinary advance in the treatment of multiple cancer types [1–4]. ICB involves the use of antibodies that target T cell co-receptors such as programmed cells death protein 1 (PD1), programmed cell death ligand 1 (PDL1) and cytotoxic T-lymphocyte associated protein 4 (CTLA-4) [5–7]. ICB has been approved for use as a first line of therapy for non-small-cell lung cancer (NSCLC), melanoma, urothelial and colorectal cancers [7]. However, only a subset of patients respond well and there is a lack of clear biomarkers that predict a favorable outcome. Some of the current predictive biomarkers for favorable ICB response involve high tumor mutation burden (TMB), increased PDL1 and PD1 expression, and tumor immune infiltration [8, 9].

Tumors with mutations in proofreading domains of DNA polymerases and mismatch repair deficiency give rise to high TMB [10, 11]. In theory, high TMB may lead to generation of mutant peptides that are processed and recognized as neoantigens by the immune system. Tumors evolve by upregulating immune checkpoints such as PD1 and PDL1, to evade the immune response and are therefore subject to favorable ICB response [12] [13]. However, some patients with high TMB still do not respond and some tumor types such as thyroid cancers with low mutation burden respond well [14]. Thus, the question of how TMB impacts tumor biology and subsequently immunotherapy response remains to be resolved. Also, there are various conflicting studies of analysis of PDL1 expression with immunohistochemistry which limit its use as a predictive marker across cancer types [15]. As ICB has been a transforming cancer treatment, there is a window of opportunity to model high TMB and ICB therapy response where the new information could be used to modify treatment protocols and utilize them for the right subset of patients.

We and others have shown that ICB therapy can have high response rates in tumors with high TMB where exceptional responses have been observed in patients with endometrial cancers, NSCLC, and colorectal cancer [9, 16–20]. NSCLC tumors with high TMB have also shown to be enriched in *Pold1* and *Pole* exonuclease mutations [18]. These exonuclease domain point mutations disable the proofreading function essential for correcting DNA replication errors, resulting in a mutator phenotype [21]. These mutations have also been associated with positive responses to ICB in several cancers [17, 22]. Tumors with somatic point mutations such as *Pole V411L* and *Pole P286R* have been shown to be associated with increased mutation burden or an ultra-mutator phenotype [17, 23–25]. In addition, the increased expression of immune checkpoint genes are also associated with this mutator phenotype making these tumors a suitable target for immunotherapy. There are other known clinically relevant germline point mutations e.g. *Pole L424V* which have been shown to be associated with predisposition to multiple cancers and an increased mutation burden [26, 27].

Traditionally, genetically engineered mouse models of cancer have been used for studying various aspects of tumorigenesis. These oncogenic driver mutation models (e.g. *Kras*-driven lung cancer models) do not present with high TMB that are exhibited by human cancers making them unsuitable for tumor-immune interaction studies [28]. Interestingly, these models also show minimal responses to ICB [29]. Therefore, there is a need for appropriate model systems that correctly represent the TMB seen in human cancers as mutational patterns both clonal and sub-clonal have shown to impact tumor biology as well as responses to chemotherapy and immunotherapy [30]. Given this background, we set out to create genetically engineered mouse models that recapitulate high mutation burdens that are inherent to many human cancers and test whether these models also respond favorably to ICB. To achieve this, we introduced point mutations in the *Pold1* and *Pole* exonuclease domains of C57BL6 mice.

The *Pold1* and *Pole* replication polymerases are responsible for lagging and leading strand synthesis, respectively [31]. Loss of exonuclease activity leads to increased errors during replication resulting in increased number of mutations during each cell division [32, 33]. We created mice models with germline *Pold1 D400A, Pole D272A E274A, Pole L424V* and conditional *Pole V411L* point mutations. These mice develop a spectrum of spontaneous tumors mainly thymic and splenic lymphomas, lung, and intestinal tumors consistent with previously published *Pold1* and *Pole* mutant models [34–36]. Exome sequencing of these tumors from these mouse models revealed a highly variable but elevated TMB with mutational signatures similar to that seen in *Pold1* and *Pole* mutant human cancers. Treatment of these elevated mutation burden mouse models with ICB caused regression of only some tumors that was insufficient to provide a survival advantage. In addition, we also bred *Pold1* and *Pole* mutant mice with LSL-*Kras G12D/+ p53 flox/flox* conditional model for NSCLC, which interestingly did not show a particularly high TMB. However, single cell passaging of tumor derived cell lines from these models modestly increased mutation burden and immunogenicity, leading to tumor immune rejection. These findings are indicative of the generation of immunogenic mutations with efficient immune editing during tumor evolution *in vivo* that is bypassed by introducing mutations *in vitro*. Finally, in contrast to treating mutator mice with ICB once tumors arose that was ineffective, treating mutator mice with ICB earlier and prior to the appearance of obvious tumors delayed cancer onset leading to improved survival, suggesting the use of ICB in high-risk settings for cancer prevention.

## Results

### Germline and conditional mouse models with *Pold1* and *Pole* exonuclease mutations have elevated mutation burden and develop spontaneous cancers

Germline *Pold1 D400A, Pole D272A E274A, Pole L424V* and Conditional *Pole V411L/+* point mutations reside in exonuclease domains of *Pold1* and *Pole* (Figure 1A) and mouse models with these mutations were created using a knock-in CRISPR-Cas9 system. We also created a conditional mouse model for *Pole V411L*, which is a commonly observed pathogenic somatic mutation associated mainly with endometrial carcinomas in humans [17]. *Pold1 D400A, Pole D272A E274A* were previously published [34, 37] but these models were not studied in the context of immunotherapy response and are unfortunately no longer available. For germline models, heterozygous cohorts of mice were bred to generate homozygous animals. The homozygous breeders were then used to generate experimental mice. For survival analysis, wild type, heterozygous and homozygous progeny from the heterozygous breeding were monitored for their lifespan. The progeny from the homozygous animals were not used further for breeding to avoid the accumulation of mutations over generations. The mice were genotyped, and sequencing analysis was done to confirm the presence of point mutations (Supplementary Figure 1A-C). The conditional *LSL-Pole V411L/+* mouse model was bred with *Ubc-creERT2/+* mice to generate *Ubc-creERT2/+; LSL-Pole V411L/+* mice where the whole-body *Pole V411L* allele is induced after tamoxifen injection (Supplementary Figure 1D). Mice were genotyped again to confirm the induction of the *Pole V411L* allele, and it was verified to be induced throughout the whole body (Supplementary Figure 1E).

**Figure 1.**
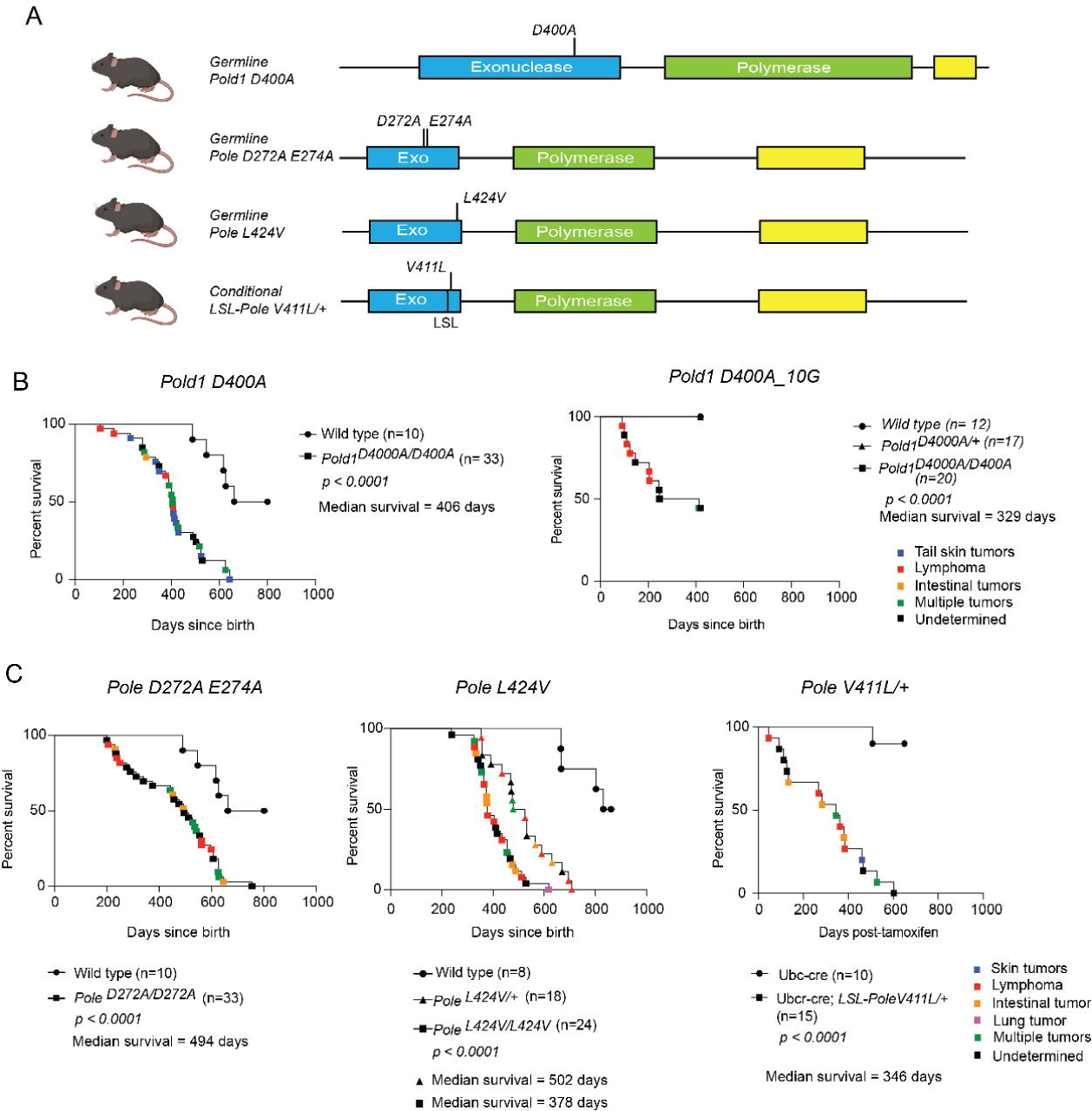
*Pold1* and *Pole* homozygous mutant mice develop spontaneous tumors and other cancers. (A) Schematic representation of mouse *Pold1* and *Pole* genes with exonuclease (EXO) and Polymerase domains with locations of mutations germline *Pold1 D400A, Pole D272A E274A, Pole L424V* and conditional *Pole V411L* with an LSL cassette. (B) Kaplan-Meier Survival curve analyses of old generation *Pold1 D400A* and new generation *Pold1 D400A_10G* mutant mice. The color key represents types of tumors detected. *p* value calculated using log rank test. (C) Survival analyses of germline *Pole D272A E274A, Pole L424V* and conditional *Pole V411L* mice. The *Pole L424V/+* showed a cancer phenotype like *Pole L424V* homozygous mutant mice. The survival analysis of *Pole V411L/+* is plotted as survival percentage against days elapsed post tamoxifen injections. The color key represents types of tumors detected. *p* value calculated using log rank test.

Both *Pold1* and *Pole* exonuclease domain mutant mice, with increased error prone mutagenesis, develop spontaneous malignancies and die of cancer. The majority (∼70%) of the spontaneous tumors seen in these models were thymic and splenic lymphomas, which often infiltrated the lungs and liver followed by intestinal and lung tumors and few cases of pancreatic tumors and sarcomas. 75% mice from the first cohort of survival analysis of *Pold1 D400A* homozygous mice presented with tail skin tumors, which were rarely seen in the *Pole* models (Figure 1B, Supplementary Figure 1G). A change in survival profile and tumor incidence was observed in later generations, in *Pold1* mice as they were bred consistently for ∼10 generations (Figure 1B). These *Pold1 D400A_10G* mice that are approximately 10 generations from the original *Pold1 D400A* mice had earlier onset of tumors, with majority of these early tumors being thymic lymphomas that reduced survival (Figure 1B). Comparison of exome sequencing of DNA from normal tissue from *Pold1 D400A_10G* mice and their ancestral *Pold1* mice revealed 113 additional mostly missense mutations and 14 genes with truncating mutations in the germline (Supplementary Table 1A). Three COSMIC cancer-associated genes (*Alk, Kmt2d,* and *Spen*) have missense mutations with *Spen* having dense missense mutations on the same allele (Supplementary Table 1B). Thus, the accelerated cancer onset may be attributed to genetic anticipation observed in *Pold1 D400A* mutant mice that were bred over time, which was characterized by acquisition of new cancer-related mutations, reduced latency, and early appearance of thymic lymphomas with successive generations.

The heterozygous *Pold1 D400A/+* and *Pole D272A/+* mice rarely showed a cancer phenotype, and their lifespan was comparable to wild type mice (Figure 1B and Supplementary Figure 1F). *Pole D272A E274A* homozygous mutant mice showed a median survival of 494 days and died predominantly from lymphomas (Figure 1C). The *Pole L424V* point mutation has been found both as a germline as well as somatic mutation in patients [26, 38]. Both heterozygous and homozygous germline *Pole L424V* mutant mice died of cancer and had a median survival of 502 and 378 days, respectively (Figure 1C) (Supplementary Figure 1G). The conditional *LSL-Pole V11L/+* mice post tamoxifen injection also succumbed to several spontaneous tumors, mainly lymphomas, intestinal tumors, pancreatic tumors, and some skin tumors and had a median survival of 346 days (Figure 1C) (Supplementary Figure 1G). Thus, all models of *Pold1* and *Pole* proofreading mutations showed accelerated tumorigenesis and reduced survival. The three *Pole* proofreading mutation models showed increased but variable tumorigenic potential as observed by the differential median survival (Figure 1C).

### The mutational landscapes of *Pold1* and *Pole* mutant tumors in mice and humans are similar

To determine the mutation burden, a set of tumors arising in *Pold1* and *Pole* mutant backgrounds were collected and subject to whole exome sequencing. This included three intestinal tumors, four tail tumors and four thymic lymphomas and one lymphoma from *Pold1 D400A* homozygous mutant mice; four intestinal tumors, three lung tumors, and two lymphomas from *Pole D272A E274A* homozygous mutant mice; two intestinal tumors, two lung tumors, one lymphoma, and three sarcomas and one thymic lymphoma from *Pole L424V/L424V* mice; five intestinal tumors and one sarcoma from *Pole L424V/+* mice, and four intestinal tumors, six lymphomas, one thymic lymphoma and one pancreatic tumor from *Pole V411L/+* mice. The total mutation frequency including single nucleotide variants (SNVs) and insertions and deletions (Indels) was increased in *Pold1* and *Pole* mutant mouse tumors (Figure 2A). In *Pold1 D400A* mutant mice, tail tumors had the highest mutation burden with over 300 mutations per Mb of exome, followed by thymic lymphomas and intestinal tumors. *Pold1 D400A* lung tumors had the lowest mutation burden (Figure 2A). In *Pole D272A E274A* mutant mice, intestinal tumors and lymphomas had the highest mutation burdens followed by lung tumors (Figure 2A). In *Pole L424V/L424V* mutant mice, intestinal tumors, lung tumors, and lymphomas all had high mutation burdens followed by sarcomas (Figure 2A). In *Pole L424V/+* mutant mice, the intestinal tumors had the highest mutation burden followed by sarcomas. In *Pole V411L/+* mice, intestinal tumors, lymphomas, and a pancreatic tumor all had high mutation burden (Figure 2A). Thus, all four models confirm that these mutator alleles increased TMB. The vast differences in TMB across different tumor types may reflect differences in the number of DNA replication cycles the tumor-initiating cell underwent prior to clonal expansion of tumor growth and establishment of the truncal mutation burden.

**Figure 2.**
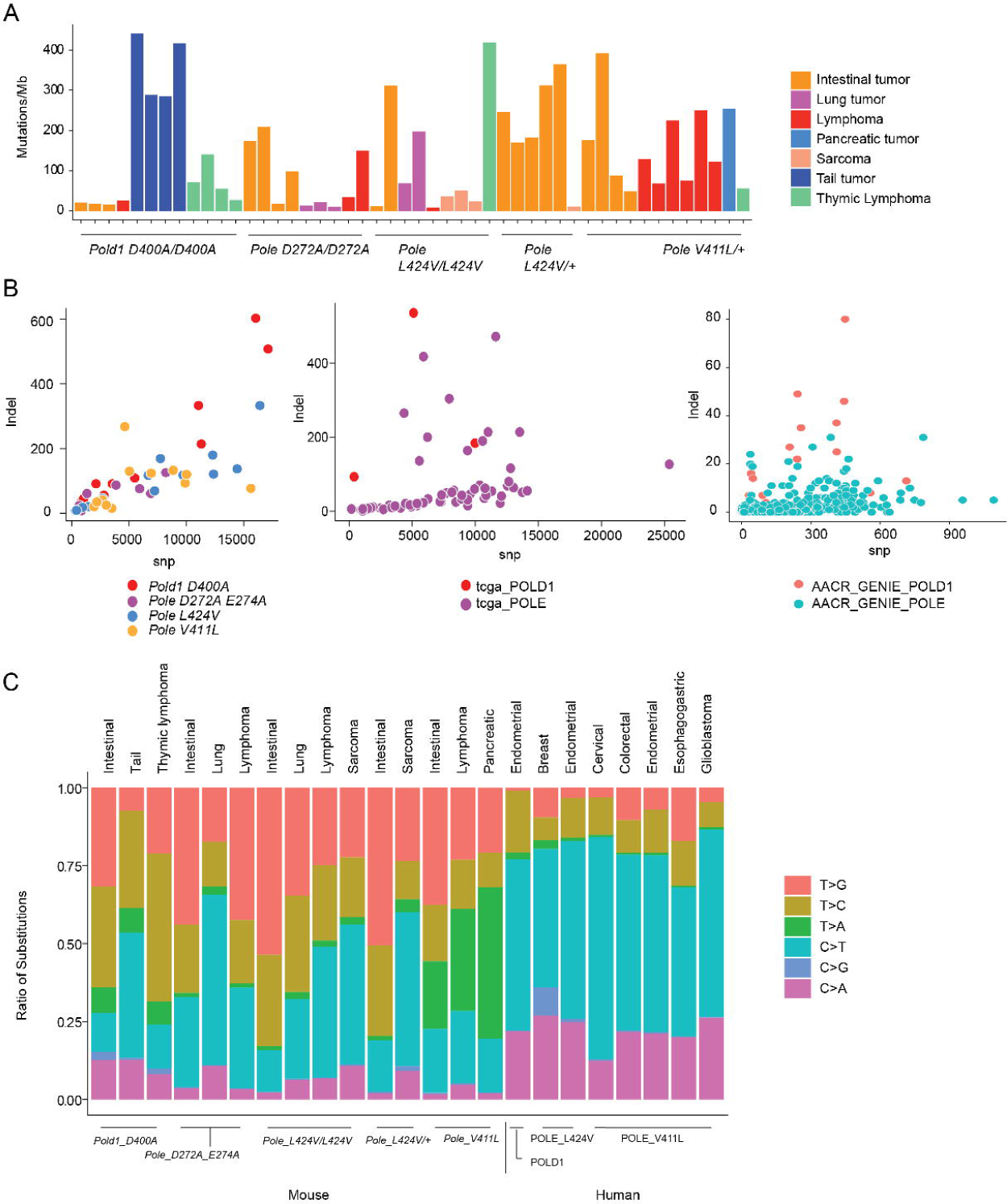
The mutational landscape of *Pold1* and *Pole* mutant tumors was similar in mice and humans. (A) Tumor mutation frequency was determined from whole exome sequencing (WES) of spontaneous tumors from *Pold1 D400A, Pole D272A E274A, Pole L424V* and *Pole V411L/+* mice. (B) Number of Indels vs SNV in spontaneous tumors from *Pold1 D400A, Pole D272A E274A, Pole L424V* and *Pole V411L/+* mice, in human *Pold1* and *Pole* mutant tumor samples from TCGA, and in human Pold1 and Pole mutant tumor samples from AACR GENIE. (C) Comparison of mutation signatures using the six-base substitution of tumor types from *Pold1 D400A, Pole D272A E274A, Pole L424V* and *Pole V411L/+* mice and in human *Pold1* and *Pole* mutant tumor samples from TCGA.

The types of mutations in the mutator mouse tumors were examined further and compared to those in humans. The relative number of Indels to SNVs is higher for tumors with *Pold1* mutations than *Pole* mutations in both mouse and human tumors from TCGA and AACR GENIE (17 *Pold1* mutant and 427 *Pole* mutant tumor samples) (Figure 2B). Similar results were observed in normal aging tissues of *Pold1* and *Pole* mutation carriers in humans [27]. These results are consistent with the role of *Pold1* in lagging strand DNA synthesis and generation of mutant Okazaki fragments that may be incorrectly processed and lead to an increased percentage of Indels. Next, the six-substitution signature analysis of TCGA and AACR-project genie cohorts confirmed similarities between the mouse and human tumor mutation signatures (Figure 2C). The COSMIC single base substitution (SBS) mutational signatures also showed similarity between the mouse and human tumors carrying *Pold1* and *Pole* mutations (Supplementary Figure 2A and 2B). Both the *Pold1* and *Pole* mutant tumors from mouse and human have SBS1 and SBS5, which are clock-like signatures as well as SBS15, which is associated with DNA mismatch repair defects. In addition, *Pole* mutant tumors also have SBS10a and SBS10b mutational signatures which were previously associated with *Pole* mutations while *Pold1* mutant tumors have SBS23 and SBS54 mutational signatures. There are differences in mutational signatures between the mice and human tumors and they may be attributed to different tumor types compared between the mice (intestinal, lymphoma, tail, lung) and human tumors (endometrial, breast, colorectal, cervical, esophagogastric, glioblastoma). The *Pole* models also showed a conspicuous presence of the SBS28 mutation signature in all tumors. The etiology of this signature is currently unknown (Supplementary Figure 2A). Thus, the mutational patterns of the polymerase mutator syndromes have similarities across mouse and humans.

Examination of the cancer-associated genes most recurrently mutated in the *Pold1* D400A mutant mice across all tumor types revealed that they were *Pten* (8/12 tumors) and *Brca2*, *Med12* and *Notch1* (7/12 tumors) (Supplementary Table 2). For tail tumors, due to the very high mutation burden, many cancer-associated genes were mutated including *Med12, Cntrl, Bub1b, Brca2, Notch1, Csmd3, Naca, Ranbp2, Fat1, Nbea, Ncor1, Spen, Fat4, Kmt2c*, and *Kmt2d* in all four tail tumors. For thymic lymphomas, the most recurrently mutated genes are *Pten, Brca2,* and *Smarca4* (3/4 tumors), consistent with loss of *Pten* being an early mutation selected in mouse thymic lymphoma development [39]. For intestinal tumors, only *Pim1* was found recurrently mutated in 2/3 tumors (Supplementary Table 2). Across all tumor types, many of the mutations in tumor suppressors (*Brca2, Csmd3, Fat1, Ncor1, Spen, Kmt2c, Kmt2d,* and *Smarca4*), are likely passenger mutations since they are heterozygous (allele frequency less than 0.5) (Supplementary Table 2A). The most recurrently mutated cancer-associated gene in *Pole* mutant mice was *Lrp1b* (25/36 tumors). For intestinal tumors, the most frequently mutated genes were *Lrp1b* (11/15), *Fat4* (11/ 15), *Muc16* (10/15), *Fas* (10/15), and *Nbea* (10/15). For lung tumors, the most frequently mutated genes were *Fat4, Csmd3, Pde4dip, Robo2, Ncor1, Med12, Grin2a, Fbxw7, Msh6*, and *Arhgef10l* (2/5 tumors). For lymphomas, the most frequently mutated genes were *Lrp1b* (8/9), *Csmd3* (6/9), and Fat3 (6/9). For sarcomas, the most frequently mutated genes were *Lrp1b, Muc16, Jak1*, and *Pten* (2/4 tumors) (Supplementary Table 3). It is intriguing that *Lrp1b* being a larger gene was found to be frequently mutated in *Pole* mutant tumors (25/36 tumors) but not in tumors from *Pold1* mutant mice (4/12 tumors) (Supplementary tables 2 and 3). However, *Lrp1b*, a purported tumor suppressor, along with tumor suppressors *Ncor1, Grin2a, Fbxw7*, and *Msh6* are likely passenger mutations since they are heterozygous (allele frequency less than 0.5) (Supplementary Table 3A). Thus, genomic analysis demonstrated that *Pold1* and *Pole* mutant tumors evolve by accumulating mutations in cancer associated genes.

### Partial response to acute ICB in *Pold1* tumors

Next, we studied the response of the *Pold1 D400A* mutant mouse tumors to acute ICB as these mice develop tumors that are visible and that have a very high TMB at ∼ 1 year of age. The *Pold1* mutant mice with apparent tail skin tumors were treated with either control IgG or Anti-PD1 antibody every five days and tumor size was monitored (Figure 3A and Supplementary Figure 3). The red arrows indicate the tumors that are shrinking, and green arrows indicate the tumors that have either increased in size or remained stable (Figure 3A). ICB has shown exceptional responses in some high TMB cancers. Anti-PD1 mediated inhibition leads to reduced immune suppression and activation and proliferation of T cells [40]. Anti-PD1 treatment of *Pold1 D400A* homozygous mutant mice showed a reduction in tumor size compared to control IgG in a subset of tumors, whereas others continued to grow or remained stable (Figure 3A and 3B). Tumors with a consistent decrease in size during the treatment period were considered as responding tumors. Anti-PD1 antibody treatment increased the percentage of responding tumors from 2.7% to 32.5%. (Figure 3C). Next, we checked for immune infiltration in the tumor sections from these mice. Immunohistochemistry analysis showed increased CD3 staining in anti-PD1 antibody-treated tumor sections (mean 6.4% positive cells) compared to Control antibody-treated group (mean 1.7% positive cells) consistent with inhibition of PD1 signaling and increased T cell proliferation (Figure 3D). However, when survival analysis was examined, there was not a significant difference between Control and Anti-PD1 antibody-treated groups, where all the mice succumbed to other cancers mainly splenic lymphomas (Figure 3E). It is possible that the lymphomas from *Pold1* mice were unresponsive to ICB because of their lower mutation burden compared to tail tumors. Alternatively, ICB may be ineffective when only a subset of tumors are responsive independent of TMB and/or when tumor burden is high along with elevation of intrinsic resistance mechanisms. These results suggested that acute ICB treatment when started after the appearance of tumors in the mutator mice increased responsiveness ten-fold but as this represented only 1/3 of the tumors it was insufficient to affect overall survival.

**Figure 3.**
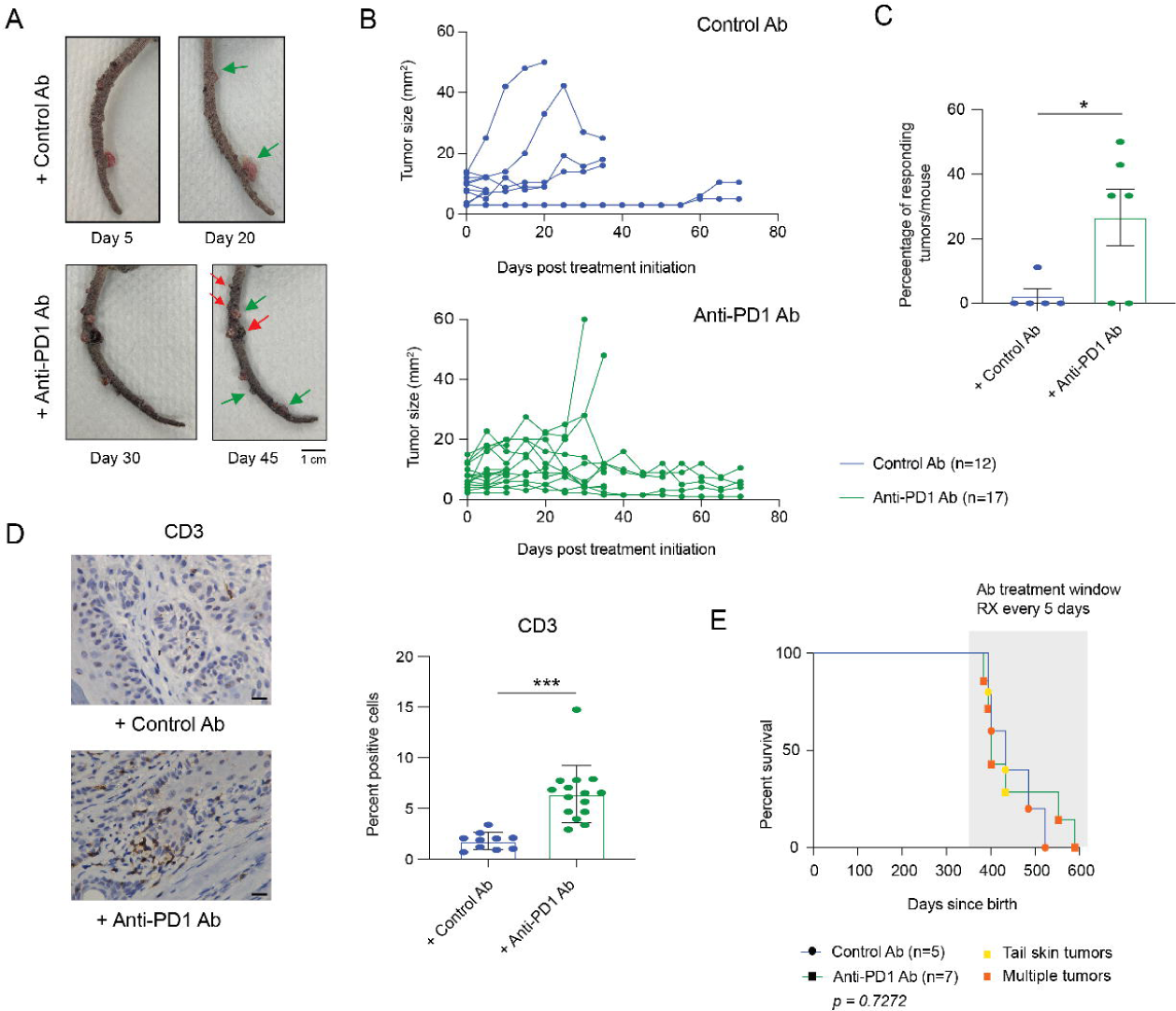
Acute ICB treatment in *Pold1* mutant mice tail tumors showed a favorable response in some tumors. (A) Gross images of mice tails with tumors treated with either control IgG or Anti-PD1 antibody. The green arrows indicate tumors that remained stable or increased in size. The red arrow indicates tumors that regressed with treatment. (B) The tumor size measurements taken and calculated every 5 days over the course of treatment for control IgG (n=12 tumors) and Anti-PD1Antibody treatment (n=17 tumors) groups of n=5 mice each. (C) Percentage of responding tumors was calculated in control IgG (n=5) and Anti-PD1 antibody (n=5) treated mice. *p <0.05 is calculated with Student’s t test. (D) Representative images of IHC for CD3 in tumor sections treated with either control IgG or Anti-PD1 antibody. The percent positive cells are plotted for tumor sections of control IgG (n=10) and Anti-PD1Antibody treatment (n=14) groups. ***p <0.001 is calculated with Student’s t test. (E) Survival analyses of Pold1 homozygous mutant mice with visible tail tumors treated with either control IgG or Anti-PD1 antibody. The light grey rectangle represents treatment window where treatment was started as the tumors appeared and continued for every 5 days for their lifetime. p > 0.05 was considered NS and calculated with log rank test. The color key represents types of tumors detected.

### Introducing *Pold1* and *Pole* proofreading mutations in autochthonous mouse models for lung cancer only moderately increased the TMB

Genetically engineered mouse models have been routinely used in studies of lung cancer tumorigenesis. Although these oncogene activation- and tumor suppressor gene inactivation-driven models have been particularly useful as they recapitulate histological characteristics of human NSCLC, they exhibit a low TMB having bypassed the normal mutagenic mechanisms that normally lead to oncogenic mutations and that generate the truncal TMB making them unsuitable for immunotherapy studies [28]. We asked whether introducing *Pold1* and *Pole* mutator alleles in these models would increase TMB and consequently modify these models to be better suitable for studying immunotherapy responses. We utilized *LSL-Kras ^G12D/+^, p53^flox/flox^*(KP) model for NSCLC [41] where the oncogenic *Kras^G12D/+^* allele is activated and *Trp53* tumor suppressor gene is deleted, mediated by intranasal delivery of adeno-Cre virus. This results in sporadic induction of oncogenic driver mutations in the lung and spontaneous lung tumor development. We bred KP mice with germline homozygous *Pold1 D400A, Pole D272A E274A, Pole L424V,* and conditional *LSL-Pole V411L/+* mice and aimed to determine the differences in low TMB (KP model) and potentially high TMB *Pol* mutator KP mice (Figure 4A). At the time of Cre-mediated tumor initiation the *Pold1 D400A*, *Pole D272A E274A* and *Pole L424V* mutations were germline, whereas the conditional *Pole V411L/+* allele was induced concurrently with the activation of *Kras^G12D/+^* and deletion of *Trp53* alleles.

**Figure 4.**
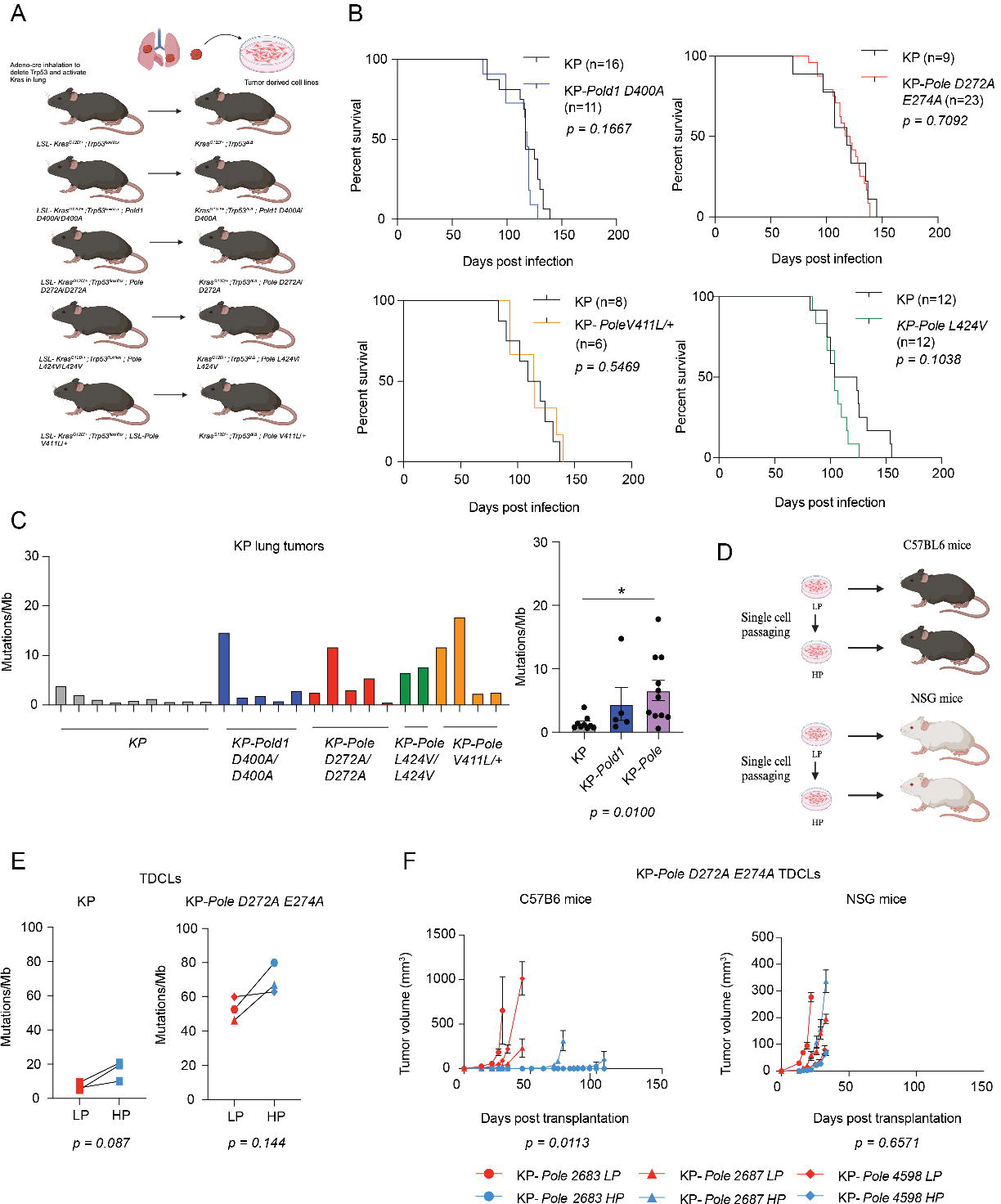
*Pold1* and *Pole* exonuclease mutations in autochthonous NSCLC mouse models moderately increased the TMB. (A) Schematic representation of the experiment where mice were nasally infected with adeno-Cre resulting in lung tumor formation. The lung tumors were then harvested to generate tumor derived cell lines (TDCLs) (created using BioRender). (B) Kaplan-Meier survival curve analyses of KP Vs KP-*Pole D272A E274A,* KP Vs KP-*Pold1 D400A*, KP Vs KP-*PoleL424V* and KP Vs KP-*PoleV411L/+* mice infected intra nasally with adeno-cre. *p* values are determined using log-rank test. (C) Total mutation frequency from WES of lung tumors from KP, KP-*Pold1 D400A* and KP-*Pole D272A E274A,* KP-*PoleL424V* and KP-*PoleV411L/+* mice at 12-week timepoint (D) Schematic representation of single cell passaged LP and HP TDCLs and subcutaneous injection in C57BL6 and NSG mice (created using BioRender). (E) Paired analysis of total mutation burden per Mb of exome in LP and HP of KP and KP-*Pole D272A E274A* cell lines. (F) Tumor growth analysis of KP-*Pole D272A E274A* low (red) and high (Sky blue) passage cell lines in immunocompetent C57BL6 and immunocompromised NSG mice. *p* values are determined using unpaired Student’s T test.

Introduction of *Pold1* and *Pole* proofreading mutations in KP mice did not affect the survival of these mice (Figure 4B) and no change in lung wet weights and immune infiltration was observed when the mice were euthanized at 12-week timepoint (Supplementary Figure 4A and 4B). These data suggested that *Pold1* and *Pole* mutator alleles did not affect tumorigenesis in KP mice. Exome sequencing analysis of lung tumors from KP and KP-*Pold1 D400A*, KP-*Pole D272A E274A*, KP*-Pole L424V* and KP*-Pole V411L/+* mice revealed only moderately increased number of mutations and did not signify a high TMB phenotype that is presented by human NSCLC cases, which have >10 mutations/Mb (Figure 4C). While the TMB is significantly higher in KP-*Pold1* and KP-*Pole* mice lung tumors compared to KP mice with wild type *Pol* alleles, the increased TMB was modest and much smaller than the spontaneous tumors that developed in the *Pold1* and *Pole* mutator mice models.

Based on the low TMB in the KP tumors with the mutator alleles, we questioned whether lack of high TMB might be the result of immune editing and elimination of immunogenic mutations occurring at a rate sufficient to permit tumorigenesis unobstructed. To address this, we created tumor-derived cell lines (TDCLs) from lung tumors of KP-Pole *D272A E274A* mice (Figure 4D). To increase the mutation burden, we passaged these cells lines as single-cell clones for up to 14 generations, which increased the mutation burden from an average of 53 exonic mutations/Mb in low passage cells (LP) cells to an average of 70 mutations/Mb in high passage (HP) cells (Figure 4E). Expectedly, the passaging of KP cells did not accumulate an increased number of mutations (Figure 4E). The LP and HP cell lines grew at similar rates *in vitro* (Supplementary Figure 4C). The karyotype and copy number profile of both KP and KP-*Pole D272A E274A* cell lines also changed from LP to HP cells (Supplementary Figure 4D). The low and high passage KP-*Pole D272A E274A* cell lines were then subcutaneously transplanted in immunocompetent (C57BL6) and immunocompromised NSG mice. We observed that LP KP-Pole *D272A E274A TDCLs* readily formed tumors in immunocompetent mice, but the HP KP-Pole *D272A E274A* TDCLs showed delayed or no tumor growth in immunocompetent mice (Figure 4F). In contrast, both LP and HP TDCLs readily formed tumors in immunocompromised NSG mice (Figure 4F). These results suggest that passaging of KP-Pole *D272A E274A* TDCLs *in vitro* where immune editing did not occur was sufficient to increase the immunogenicity for tumor rejection by the immune system. Altogether, these results suggested that introducing *Pold1* and *Pole* mutator alleles in autochthonous mouse models of lung cancer only moderately increased the mutation burden. However, single-cell passaged *KP-Pole D272A* TDCLs were able to accumulate additional mutations resulting in increased immunogenicity and tumor rejection suggesting mechanisms of immunoediting are at play *in vivo*.

### Prophylactic ICB delays cancer onset and improves survival of mice with *Pold1* and *Pole* proofreading mutations

As acute ICB treatment of *Pold1 D400A* homozygous mutant mice with overt cancer showed only a partial response in a subset of tumors and no survival advantage, we tested whether an earlier intervention where tumor numbers, heterogeneity, and burden are reduced, could improve response. ICB was started when the mice reached adult age at 8-10 weeks old and prior to the appearance of overt cancer, and their survival and tumor profile were monitored. Two clones of anti-PD1 antibodies-RMP-14 and 29F.1A12 clones were used for biweekly treatments in all *Pold1* and *Pole* proofreading mutation mouse models. The mouse PD-1 cDNA was used as an immunogen to produce RMP-14, whereas a recombinant PD-1-Ig fusion protein was used to produce 29F.1A12. *In vitro* binding assays using these 2 clones of antibodies have shown differences in avidity and epitopes recognized [42]. The treatment frequency for the cancer prevention strategy was reduced to biweekly treatments compared to the acute treatment of existing tumors (treatment every 5 days) in anticipation that less dosing may be effective with reduced tumor burden, heterogeneity, and progression. The anti-PD1 RMP-14 treatment in *Pold1 D400A* homozygous mutant mice improved survival compared to control IgG treated group (*p= 0.0388* Grehan-Breslow Wilcoxon test) (Figure 5A). The median survival for control IgG and anti-PD1 treated groups showed improvement from 223 to 349 days (Supplementary Figure 5A). The cause of death was malignancies, mainly lymphomas, which were seen earlier, and intestinal tumors and tail tumors which were seen as the mice got older (Supplementary Figure 5A). Next, whole exome sequencing of the tumors that appeared later was performed to check for any mutations that are associated with immunotherapy resistance mechanisms. Mutational analysis of the control IgG and anti-PD1 antibody-treated group of tumors showed increased TMB for tail tumors but decreased TMB for intestinal and thymic lymphoma in the treated group (Figure 5B). The most recurrently mutated COSMIC cancer gene in Anti-PD1 treated tumors is *Gnas* (4/6 tumors) while in control antibody treated tumors is *Pten* (3/6 tumors) (Supplementary tables 4 and 5) and have been previously implicated in modulating ICB response [43, 44]. Also, all six of the anti-PD1 treated tumors were diploid while the control antibody treated tumors showed a trend of increased amount of CNVs compared to anti-PD1 treated tumors and 2/6 tumors displayed high level of aneuploidy (Figure 5C and 5D). Recurrent CNVs exclusive to Anti-PD1 treated tumors were identified. (Supplementary Table 6). Of note, the *Apol7c* locus is significantly amplified in three anti-PD1 treated tumors but not in any control antibody treated tumors and has been found to play a role in antigen cross-presentation by dendritic cells [45]. These results suggest that anti-PD1 treatment may have stronger selection for clones without aneuploidy and fewer number of CNVs than in control antibody treated tumors. The same experiment was performed with 29F1.A12 clone of anti-PD1 antibody, however, a significant improvement in survival as was seen with the RMP-14 clone was not observed (Figure 5A, Supplementary Figure 5B). These findings suggest that ICB can have improved efficacy when used early, prior to the obvious appearance of tumors. Why one ICB clone would have superior efficacy than another has been reported, but the mechanism is unclear [42].

**Figure 5.**
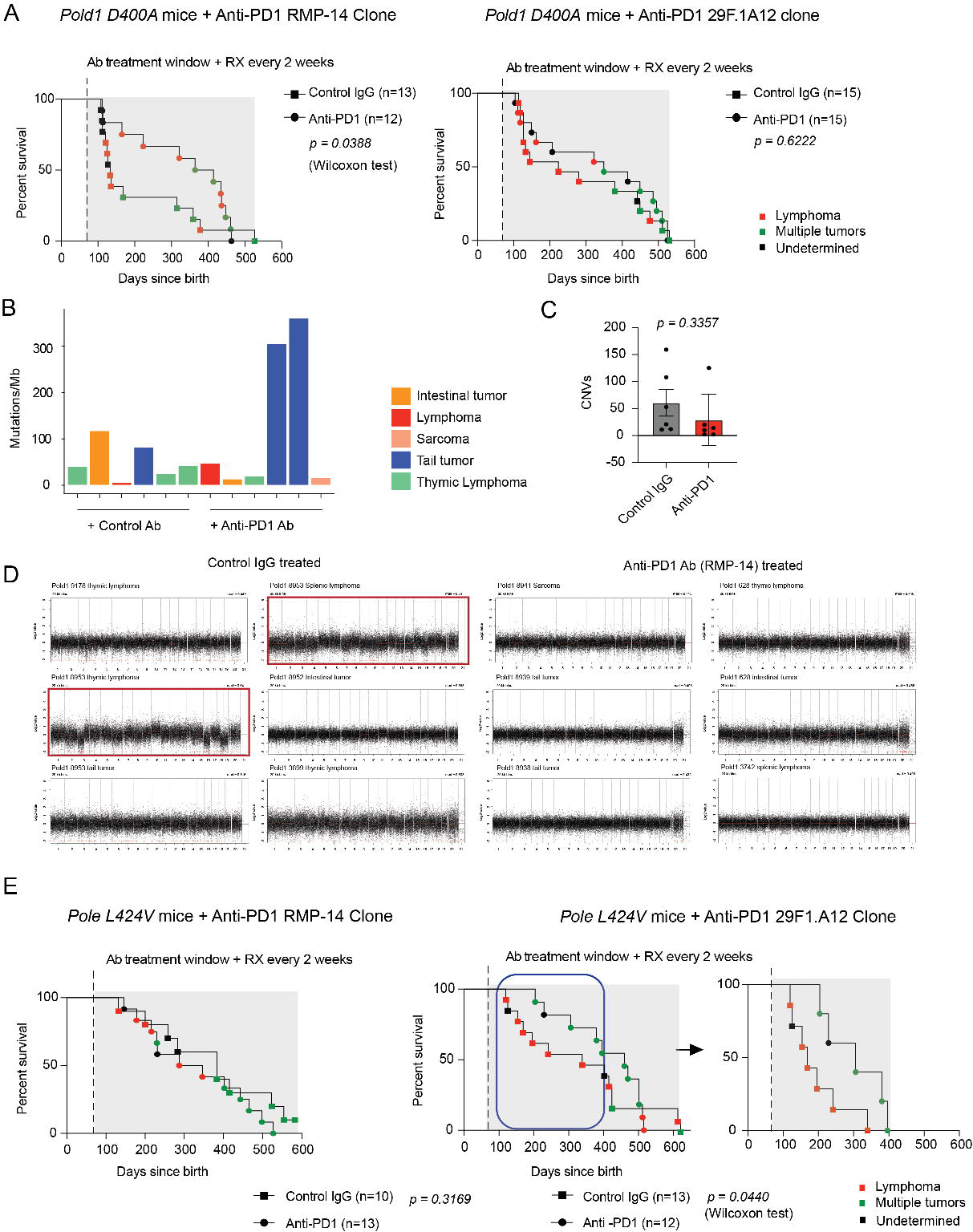
Prophylactic ICB treatment delayed cancer onset in *Pold1* and *Pole* homozygous mutant mice. (A) Survival analyses of 8–10-week-old Pold1 D400A homozygous mutant mice treated with either control IgG or Anti-PD1 antibody (RMP-14 clone) * p < 0.05 Grehan Breslow test and (29F.1A12 clone) (p = NS) with their respective control IgG antibodies. The color key represents types of tumors detected. The light grey rectangle represents the treatment window where treatment was started when the mice reached adult age of 8–10-week-old and continued every 2 weeks for their lifespan. (B) Total mutation frequency from WES of control ab and anti-PD1 ab (RMP-14 clone) treated *Pold1 D400A* tumors. (C) The quantification of CNVs in control antibody (n=6) and anti-PD1 antibody treated tumors (n=6) (D) CNV profiles of tumors from Control IgG treated and Anti-PD1 Ab RMP-14) treated mice. The samples with high level of aneuploidy are highlighted with red rectangles. (E) Survival analyses of 8–10-week-old Pole L424V homozygous mutant mice treated with either control IgG or Anti-PD1 antibody (RMP-14 clone) p =NS and (29F.1A12 clone) (p = 0.0440 Grehan-Breslow Wilcoxon test) with their respective control IgG antibodies. The color key represents types of tumors detected.

Next, we performed similar experiments with these 2 clones of ICB antibodies in the *Pole L424V* mouse model. This germline mouse model is an ideal test group for these preventive studies as this exact mutation is associated with a predisposition to multiple cancers including polyposis and colorectal cancers in humans [46]. We used homozygous mutant mice at 8-10 weeks old age and continued biweekly treatments with either control IgG or anti-PD1 antibody clones (Figure 5E). In contrast to the *Pold1 D400A* mouse model, we observed that anti-PD1 antibody treatment with the 29F.1A12 clone in *Pole L424V* mutant mice initially delayed cancer onset and improved survival (*p = 0.0440* Grehan-Breslow Wilcoxon Test) compared to the RMP-14 clone treated cohort (Figure 5B). The mice died of mostly lymphomas and intestinal tumors in these experiments (Supplementary Figures 5C and 5D). These findings support the superior efficacy of earlier intervention with ICB in cancers produced by polymerase mutator syndromes. They also demonstrate that the elevated TMB produced by polymerase mutator syndromes confers response to ICB. These mouse models provide a foundation for investigating the mechanisms controlling response to ICB and for probing the means to improve effectiveness through combinatorial approaches.

## Discussion

In this report, we established mouse models with homozygous point mutations in exonuclease domains of the replicative DNA polymerases *Pold1* and *Pole*, two of which (*Pole L424V* and *V411L*) are also seen in humans. These mutations disable the proofreading functions of the polymerase and lead to increased number of somatic mutations with each cell division. *Pold1* and *Pole* mutant mice developed spontaneous tumors, with different tissue specificity, suggesting that *Pold1* and *Pole* mutations influenced tissue-specific differential tumor development as previously observed [34]. This may be due to the specific mutational signature and its propensity to mutate oncogenic drivers in a particular tissue and the number of DNA replication cycles of the tissue stem cells or progenitors required to generate these mutations.

Patients with *Pole* mutations have been reported to be prone to colorectal and endometrial cancers [46]. Introducing *Pole* exonuclease domain mutations in C57BL6 mice caused them to mainly develop lymphomas and some intestinal and lung tumors shown in this study and others [36, 47]. The types of tumors formed appear to be influenced by the mouse background as *Pole* mutations leading to endometrial tumors with high TMB have been reported [23]. However, the mutation signatures reported in mouse models with endometrial tumors did not completely recapitulate genomic profile of human cancers with *Pole* proofreading mutations [23]. The survival analyses and tumor incidence data in the current study showed appearance of multiple tumors in *Pole* and *Pold1* mutant mice suggesting accumulation of mutations leading to high TMB and possible acquisition of new driver and passenger mutations (Supplementary Tables 2 and 3). Six-substitution analysis of the tumors from all four of our models indicated that the mutation signatures in mouse tumors were comparable to those in *Pold1* and *Pole* mutant human cancers. In addition, in the *Pold1 D400A* mouse model, tail skin tumors showed increased numbers of mutations compared to lymphomas and intestinal tumors suggesting tissue-specific differences in the rates of mutation accumulation or clonality. As mutation frequency depends on number of rounds of DNA replication the rates of stem cell divisions, and number of cell divisions required to generate a tumor-initiating clone, may be responsible for differential mutation burden detected in tumors originating from different tissues [48, 49].

*Pold1 D400A* spontaneous tumors showed an increased percentage of Indels compared to the *Pole* mutant spontaneous tumors, which is consistent with the previous data from normal and neoplastic tissue samples from patients with germline *POLD1* and *POLE* mutations [27]. These results can be explained by the *Pole* being involved in the DNA replication of the leading strand while *Pold1* is involved with the lagging strand where DNA mismatches induced by mutant *POLD1* may impair effective ligation of Okazaki fragments [50–52]. This increased number of Indels over SNPs may also contribute towards increasing immunogenicity in *Pold1 D400A* mice and partial favorable response to immunotherapy [53]. Whether the increased percentage of indels in *Pold1* mutant mice compared to *Pole* mice also contribute towards differences in neoepitopes generated and consequently the favorable clone of anti-PD1 antibody that worked remains to be understood.

It is interesting to note that introducing *Pold1* and *Pole* proofreading mutations in the KP NSCLC mouse model only moderately increased the mutation burden and did not result in more immunogenic lung tumors. However, single cell passaging of TDCLs from these tumors showed increased immunogenicity where the high passage cells were rejected in immunocompetent but not immunodeficient hosts. These results point to the possibility of immune editing *in vivo* where there is loss of neoantigens due to immunoselection where the cells continue to accumulate mutations in the absence of the immune system *in vitro* [54]. Furthermore, the presence of germline *Pold1 D400A, Pole D272A; E274A* and *Pole L424V* mutations may develop tolerogenic immune mechanisms that suppress immune response to these “self-antigens” present since birth. The conditional LSL-*Pole* V411L/+ allele when induced concurrently with *Kras* activation and *p53* deletion only moderately increased the TMB suggesting that the accelerated tumor growth rates and loss of *p53* and subsequent loss of antigen presentation may further support immune suppressive mechanisms [55]. A recent study showed that mismatch repair deficiency when introduced simultaneously with oncogenic mutations, did not greatly increase clonal TMB or provide any therapeutic benefit with ICB and suggested that increased intratumor heterogeneity is responsible for poor neoantigen expression and reduced immune infiltration [56]. However, timing of MMR defect may also be critical to increase TMB, as the MMR defect may need to occur well before accumulation/introduction of oncogenic mutations to allow increase of truncal mutation burden, and thus result in high truncal mutation burden after oncogenic transformation in the subsequent cancers. Low neoantigen expression was also shown to be responsible for T cell dysfunction and immune evasion in a colorectal cancer model [57]. These new data suggest that there might be factors other than TMB alone that modulate the efficacy of ICB. It is still remarkable how efficiently mutator tumors overcome the immune response against them through their ability to acquire advantageous mutation while also how little impact the occurrence of immunogenic mutations matters once tumors progress.

An intriguing finding from our study is that tumors from control antibody treated mice showed increased CNVs and aneuploidy, whereas the anti-PD1 antibody treated tumors were all diploid. Recent studies have shown that highly aneuploid tumors are associated with markers of immune evasion and respond poorly to immunotherapy [58, 59]. Whether there is immune selection pressure and loss of heterogenous populations of aneuploid cells in the presence of continuous anti-PD1 treatment remains to be determined. In addition, the WES analysis of tumors from control and anti-PD1 antibody groups also showed the amplified *Apo17c* locus in anti-PD1 treated tumors. It is not clear whether this genomic alteration is associated with any possible resistance mechanisms involving antigen presentation pathways. With anti-PD1 treatments, there is initial improvement in tumor free survival but the tumors appearing later may be developing ICB resistance. It will be interesting to study whether a combination of anti-PD1 and anti-CTLA4 treatment shows any improved survival as this may target these co-inhibitory receptors in early priming phase as well.

With this study we conclude that exonuclease domain point mutations in *Pold1* and *Pole* generated spontaneous tumors with high mutation burden and recapitulated genomic profiles of human cancers where ICB response was improved by treating the mutator mice earlier. Thus, ICB is more effective with less tumor burden and earlier in tumor evolution suggesting a prevention approach for use of ICB for individuals with a known cancer risk. These models will also be beneficial for testing future immune oncology drugs and combination therapies as tumors developed spontaneously with increased accumulation of mutations overtime and mimicked genomic features of their human counterparts.

## Materials and Methods

### Mice

All mice were maintained and bred in compliance with the Rutgers University Institutional Animal Care and Use Committee (IACUC) guidelines. *Pold1* D400A mutant mice were created by electroporation of C57BL6/J zygotes with a mixture of 50ng/uL Cas9 protein (Millipore Sigma), .6pmol/uL each crRNA (spacer sequence CCCGAGAGATGAGGTATGGG; NCBI GRCm38.4 Chr7:44540539-44540558(+)) and tracrRNA (Millipore Sigma) and 50ng/uL ssODN donor (5’TCTGAAGAGAAGGTAGCTGAGGTAAAGACAGCCTGGTTGGCCTTTGTGTCCGTCCCCTCCACAGGCCTGGGCCGACTTCATCCTTGCCATGGACCCTGACGTGATCACCGGCTACAACATTCAGAACTTTGcaCTCCCATACCTCATCTCTCGGGCACAGGCCCTAAAGGTGAGGGAAGC, lower case is sequence change) (IDT, Ultramer). Intact electroporated zygotes were transferred on the same day into pseudo pregnant mice and allowed to give birth to potential founders. 4 founder mice had the D400A change determined by Sanger sequence analysis. PCR primers to amplify the region are POLDA 5’TTGCAAGTGCGGAGGTTGTCTTGG and POLDB 5’CTGTCCACAGCGACGAATTTCCG.

*Pole D272A E274A* mutant mice were created by electroporation of C57BL6/J zygotes with a mixture of 50ng/ul Cas9 protein (Millipore Sigma), .6pmol/ul each crRNA (spacer sequence AGGGAATTTGAGAGGCAGTT; NCBI GRCm38.4 Chr5:110294546-110294565(−)) and tracrRNA (Millipore Sigma) and 50ng/ul ssODN donor (5’GGCTGAATATGGGTGTCCAAAGGATGTATCCAGACCTGAATTCTCCGTATATTTTGTCCTTTTAGGACCCTGTGGTTTTGGCATTTGcCATCGcGACGACtAAACTGCCTCTCAAA TTCCCTGATGCTGAGACCGATCAGAT, lower case is sequence change) (IDT, Ultramer). Intact electroporated zygotes were transferred on the same day into pseudo pregnant mice and allowed to give birth to potential founders. 3 founder mice had both D272A, E274A changes determined by Sanger sequence analysis. PCR primers to amplify the region are POLEA 5’TGCGGTACTGGTGAGTGAACCTAG and POLEB 5’TCAGAAGGCAGATGCAGGAGAACC.

*Pole L424V* mutant mice were created by electroporation of C57BL6/J zygotes with a mixture of 50ng/ul Cas9 protein (Millipore Sigma), .6pmol/ul each crRNA (spacer sequence TGTGGGCAGTCATAATCTCA; NCBI GRCm39 Chr5:110444899-110444918(+)) and tracrRNA (Millipore Sigma) and 50ng/ul ssODN donor (5’GTCCTCAGGGTCCAGCTCTACAGGGTCATAGCCAAGTTTGGCCTTGGCAGCTGCtTT aAcATTATGACTGCCCACAGGAAGGTAACTGTCCCTCTTCACCCACCTGGAAAAAACA CAGATTC-3’, lower case is sequence change) (IDT, Ultramer). Intact electroporated zygotes were transferred on the same day into pseudo pregnant mice and allowed to give birth to potential founders. 1 founder with the L424V was determined by restriction digest with MseI (NEB) and Sanger sequence analysis. PCR primers to amplify the region are POLEC 5’ ACCTGAGAGTCAAGTGGAGGCTC and POLED 5’ GAAAAGCTAGTACCCATCTCAGGTAAC.

*LSL-Pole-V411L* mutant mice were generated by microinjecting into pronuclei of C57BL/6J zygotes a mixture of 50ng/ul Cas9 protein (Millipore Sigma), .6pmol/ul sgRNA (single guide RNA with spacer sequence AGTGGAGGCTCAAGTGGCAT (Millipore Sigma); NCBI GRCm39

Chr5:110444754-110444773(+)) and 10ng/ul of donor plasmid DNA. The plasmid construct consisted of 4kb 4xPolyA-stop-cassette [24] flanked with approximately 1.4Kb of homology arms on each side; however, the 3’ arm contains the modified exon13 with V411 codon replaced by CTT codon for Leucine.

Injected zygotes were transferred on the same day into pseudo pregnant mice and allowed to give birth to potential founders. 5 founder mice were found with the cassette properly targeted into the locus as determined by long PCR reactions; 3 of those founders were confirmed by Sanger sequencing to carry, along with the cassette, the V411L mutation determined by Sanger sequencing analysis. PCR primers to amplify the region encompassing the mutation are P286RC 5’AATTCCGCAAGCTAGCCACC and POLEF 5’ AGCTCTACAGGGTCATAGCCAAG.

### Tamoxifen injections

The *Pole V411L/+* allele was induced by intraperitoneal tamoxifen injections by activating Cre in *Ubc-CreERT2/+*; *LSL-PoleV411L/+* mice. Tamoxifen (T5648 Sigma) was dissolved at the concentration of 20 mg/mL in 98% sunflower seed oil and 2% ethanol. 10 uL/g body weight was intraperitoneally injected once per day for 4 consecutive days.

The mice were allowed to recover from tamoxifen treatment for a week and samples were taken for genotyping to confirm the *Pole V411L/+* allele activation. The PCR primers used to confirm activation were POLEC 5’ ACCTGAGAGTCAAGTGGAGGCTC and POLED 5’ GAAAAGCTAGTACCCATCTCAGGTAAC.

### Whole exome sequencing analysis

Fresh frozen mouse tissue for tumor and corresponding normal samples was collected at fixed or humane endpoints. The fresh frozen tissues were submitted to Azenta Life Sciences for whole exome sequencing. Samples were collected for sequencing on the Illumina platform with a 2×150 bp configuration. Sequence quality control was conducted using fastQC v0.12.1, and adapter removal as well as read trimming were performed using Trimmomatic v0.39 in paired-end mode [60]. Subsequently, sequences were aligned against the mouse reference GRCm38 using bwa-mem 0.7.17 (Li H 2013). Qualimap 2.3 [61] was used to examine sequencing alignment and facilitate the quality control of the data in SAM/BAM format; samtools (GigaScience) were used to sort and index the BAM files. Base recalibration was performed using GATK 4.2.5 [62] with known-site dataset from the Mouse Genomes Project (https://www.sanger.ac.uk/data/mouse-genomes-project/). The final BAM files were also marked as duplicated, sorted, and indexed using GATK tools. GATK Mutect2 was used for somatic mutation calling with a panel of normal and tumor normal paired mode as described previously [63, 64], and mutect-filter in GATK Mutect2 was used for an initial filter on the variants. A hard filter was also performed using bcftools v1.11-3 (GigaScience), with tumor AD (Allelic depths for the ref and alt alleles in the order listed) >=3, DP (Approximate read depth (reads with MQ=255 or with bad mates are filtered)) >=10 for tumor and normal both, and tumor AF (Allele fractions of alternate alleles in the tumor) >=0.1. Snpeff 5.2 [65] was used for annotating and predicting the effects of genetic variants on genes and proteins. To remove germline mutations artifact in this pipeline, we filtered the recurrent mutations that occurred in more than one sample within the group. Recurrently mutated genes were identified using Snpsift v5.0e [66] and Linux awk/grep commands. Python 3.9.12 and Rstudio 2022.12.0+353/R 4.2.1 were used for data visualization.

For mutational signature analysis, SNVs, insertions, and deletions were counted using Linux commands awk, sed, and grep; and bcftools v1.11-3. By considering the pyrimidines of the Watson-Crick base pairs, we counted six different possible substitutions: C>A, C>G, C>T, T>A, T>C, and T>G using Linux command awk/grep. To further characterize the mutation signatures, we used COSMIC mutation signatures v3.4 (https://cancer.sanger.ac.uk/signatures/) and SigProfiler Bioinformatic Tools, SigProfilerExtractor v1.1.23.

### De novo germline mutation calling

A cohort-based pipeline employing GATK v4.3.5 HaplotypeCaller with the GRCm38 reference was used for de-novo germline mutation calling on individuals. For cohort population integration, gatk VariantRecalibrator and applyVQSR were applied separately under SNP and INDEL modes. Private variants from distinct groups were extracted by utilizing bcftools v1.11-3. Final filters used are cohort DP > 250 and AF >= 0.4.

### Copy Number Variant analysis

The R package CopywriteR [67] v2.22.0 employing paired tumor samples with their respective normal controls was used for CNV analysis. Segmentation was performed using a 20k bin-size based on the reference GRCm38, which divides the genome into regions of consistent copy number alterations. Subsequently, CNVs were called based on those segmented data using log2-transformed ratio thresholds.

### Adeno-cre virus infection

KP Mice were anaesthetized and infected intranasally with adenovirus expressing Cre recombinase (University of Iowa Adenoviral Core) at 4 × 10^7^ plaque-forming units (pfu) per mouse when the mice were at 8-10-week-old age to induce lung tumors.

### Incucyte growth assays

∼20000 cells were plated in RPMI media supplemented with 10% Fetal Bovine serum and 1@ Penicillin-streptomycin solution in 24-well plates. The cells were allowed to grow for 4 days and monitored for percent confluence using Incucyte live cell imaging and analysis system.

### Flow cytometry

Lung tumors from KP models were harvested at 12-week timepoint and were homogenized in RPMI medium supplemented with 10% FBS (Gibco) using gentleMACS Octo disscociator (Miltenyi Biotec) according to the manufacturer’s protocol. Immune cell collection and cell surface immunostaining was performed with anti CD3 antibody (1:300 dilution, clone 17A2, 56-0032-82). Data were obtained using an LSR-II flow cytometer (BD Biosciences) and analyzed with FlowJo software (Tree star).

### Immunohistochemistry

Mouse normal and tumor tissues were fixed in 10% formalin overnight and were stored in 70% ethanol before paraffin embedding. For antibody staining, the paraffin cut tissue sections were deparaffinized with xylene and ethanol, rehydrated and were boiled for 30 minutes in 10 mM citrate buffer (pH 6.0). The sections were blocked with 10% goat serum in PBS for 1h at room temperature. The primary antibody incubation (Anti-CD3 antibody, Abcam Ab16669 1:100) was performed overnight at 4C. Next day, anti-biotinylated secondary antibody incubation was done for 20 minutes at RT followed by 3% hydrogen peroxide for 5 minutes, horseradish peroxidase streptavidin solution (SA-5704 Vector laboratories) for 15 minutes. Next, the slides were washed and developed with DAB (Vector laboratories) according to the manufacturer’s instructions. The sections were counterstained with hematoxylin, dehydrated and mounted with Cytoseal mounting media (Thermo Scientific). The images were taken at 60X using Nikon Eclipse 80i microscope and at least 10 images per condition were quantified for percent positive cells.

### ICB Antibody treatments

*Pold1* mice with tail tumors were treated with either either 250 ug of control IgG (BioXcell #BE0089) or Anti-PD1 antibody (RMP-14 clone BioXcell #BE0146) every 5 days. *Pold1* and *Pole* mutant mice treated with either 250 ug of control IgG (BioXcell #BE0089) or Anti-PD1 antibody (RMP-14 clone BioXcell #BE0146) and 200 ug of either control IgG (BioXcell #BE0089) or Anti-PD1 antibody (29F.1A12 clone BioXcell #BE0273) once every 2 weeks for their lifespan.

### Statistics

All statistical analyses were performed using Graphpad Prism v9.5.0. software. The sample size was chosen in advance depending on the available number of mice for each experiment. The mice were allocated randomly to experimental groups and no statistical methods were used to predetermine sample sizes.

## Author contributions

AS designed, performed most of the experiments and wrote the manuscript. FS performed genomic data analysis. EC assisted with KP mice experiments. ZH assisted with maintaining mouse colonies, ear tagging and antibody treatments. SA, JB assisted with genotyping, cell culture and IHCs. JP helped with ear tagging and genotyping. JDR and CSH provided valuable suggestions. CSC supervised genomic study, interpreted results, and assisted with writing. EW, ECL and SG conceived and supervised the study. All authors helped with the review and provided their comments.

## Acknowledgements

This work was supported by NIH grant R01 CA243547 awarded to EW, SG and EL, and R01 CA163591 to EW and JDR, Ludwig Institute for Cancer Research, Ludwig Princeton Branch to EW, JDR. Duncan and Nancy MacMillan Center of Excellence in Cancer Immunology and Metabolism to EW and CSH. AS was supported by postdoctoral fellowship DFHS10PPC029 from New Jersey Commission on Cancer Research (NJCCR). The authors acknowledge Drs. Peter Romanienko, Ghassan Yehia and the Rutgers Genomic Editing Shared resource for their support with creating the mouse models. The authors also thank Joshua Vieth and Shashi Sharma from the Rutgers immune monitoring shared resource for their help with flow cytometry and Dr. Jian Cao for his insights on clones of anti-PD1 antibodies. We thank Fallon Wald and Mazen Hafiz for their help with genotyping and IHC optimization. The authors acknowledge Bioinformatics Shared Resource (NCI-CCSG P30CA072720-5917). The Biospecimen Repository and Histopathology Service Shared Resource from the Rutgers Cancer Institute of New Jersey provided histology services (P30CA072720-5919). The schematic representations were created using BioRender. The authors also acknowledge the support from all members of the White laboratory.

## Supplementary Figures

**Supplementary Figure 1.**
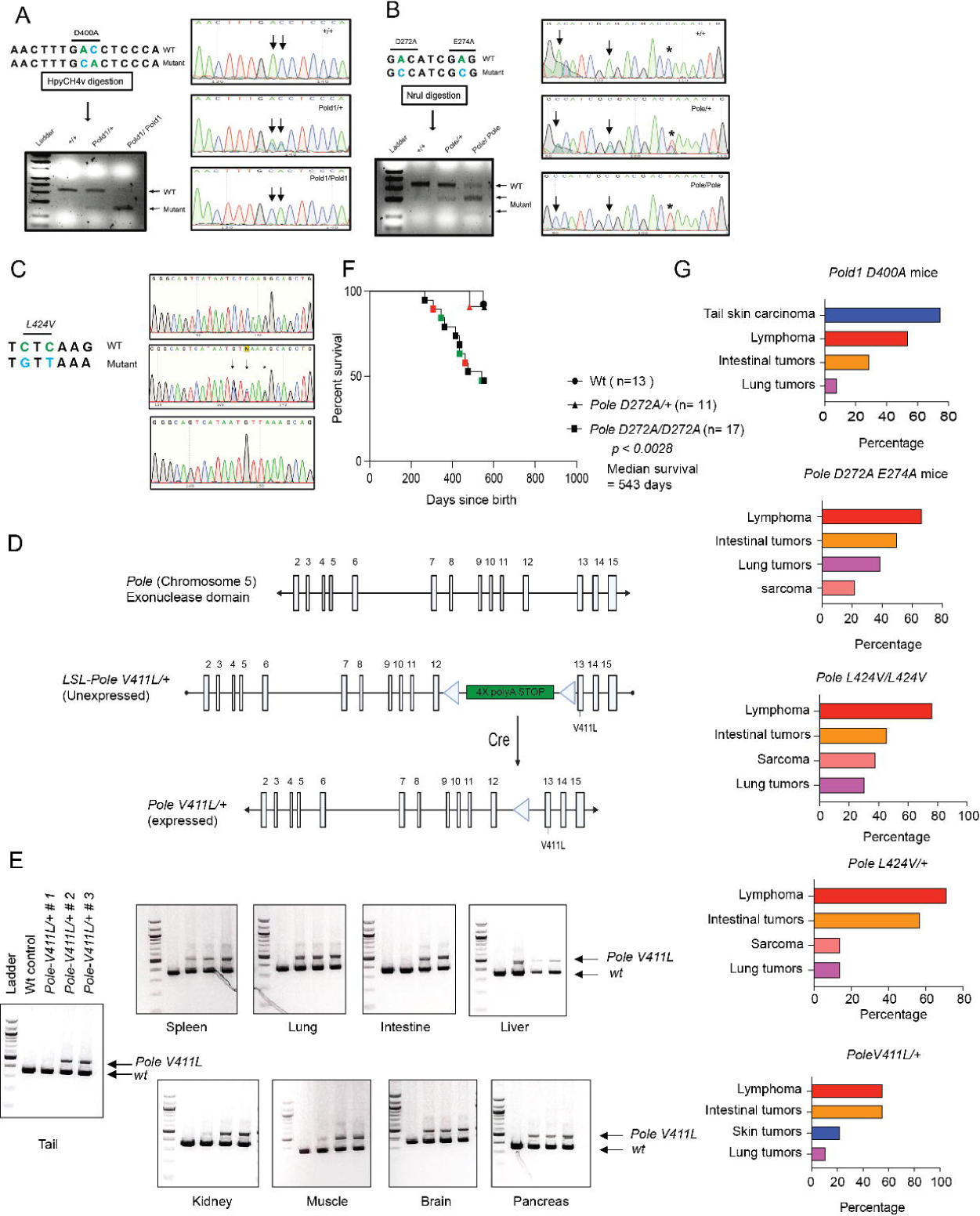
(A) Schematic representation of *Pold1 wt* sequence and sequence with point mutation *D400A*. The restriction digestion enzyme HpyCH4V was used to cut at the site of point mutation. The digested products were run on 2% agarose gels. The band sizes for wt and mutant DNA sequences are indicated for wt, heterozygous and homozygous mice. The Sanger sequencing of PCR products was carried out and trace files are shown with their indicated genotype. (B) Pole mutant sequence is represented with point mutations *D272A* and *E274A*. The restriction enzyme NruI was used to cut the mutant sequence and detect the genotype status of Pole mice. Also, sanger sequencing of PCR products was carried out and trace files are presented with their indicated genotype. * Represents silent mutation added to prevent degradation of the donor sequence by Cas9. (C) Schematic representation of *Pole* wt sequence and sequence with point mutation *L424V*. (D) Schematic representation of Pole Chr 5 exons and location of LSL casset. Post Cre activation, *Pole V411L/+* allele is induced in the whole body. (E) The DNA samples were genotyped post TAM induction and *Pole V411L/+* activation was confirmed in major organs. (F) Survival analysis of wt, heterozygous and homozygous *Pole D272A E274A* mice. (G) Tumor incidence profiles of *Pold1 D400A*, *Pole D272A E274A,* Heterozygous and homozygous *Pole L424V* and *Pole V411L/+* mice post TAM induction.

**Supplementary Figure 2.**
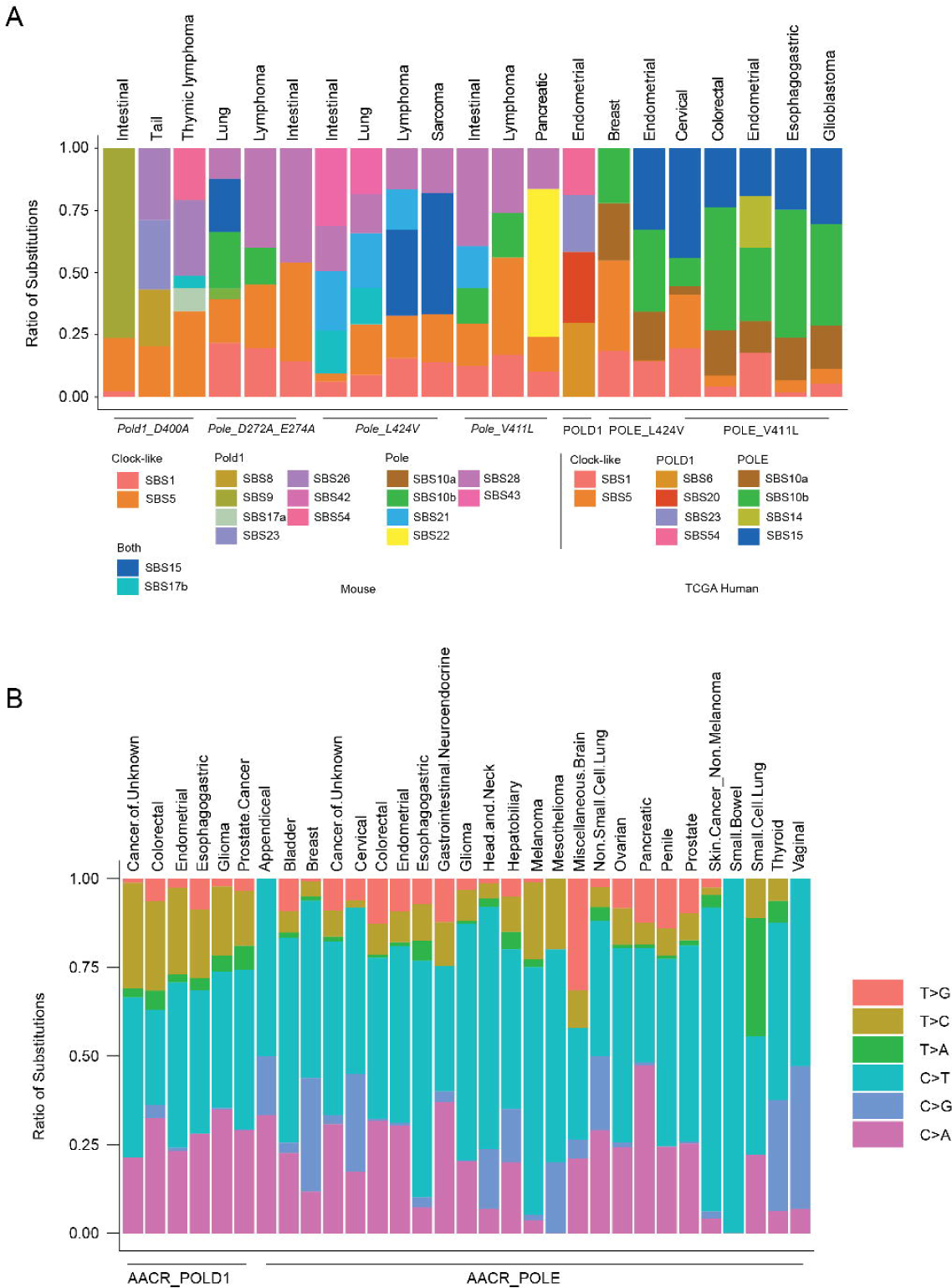
(A) COSMIC SBS mutational signatures of *Pold1* and *Pole* mutant mouse tumor types and TCGA *Pold1* and *Pole* mutant tumor types. (B) Six-substitution signature analysis in *Pold1* and *Pole* mutant human cancers from AACR Project Genie dataset.

**Supplementary Figure 3.**
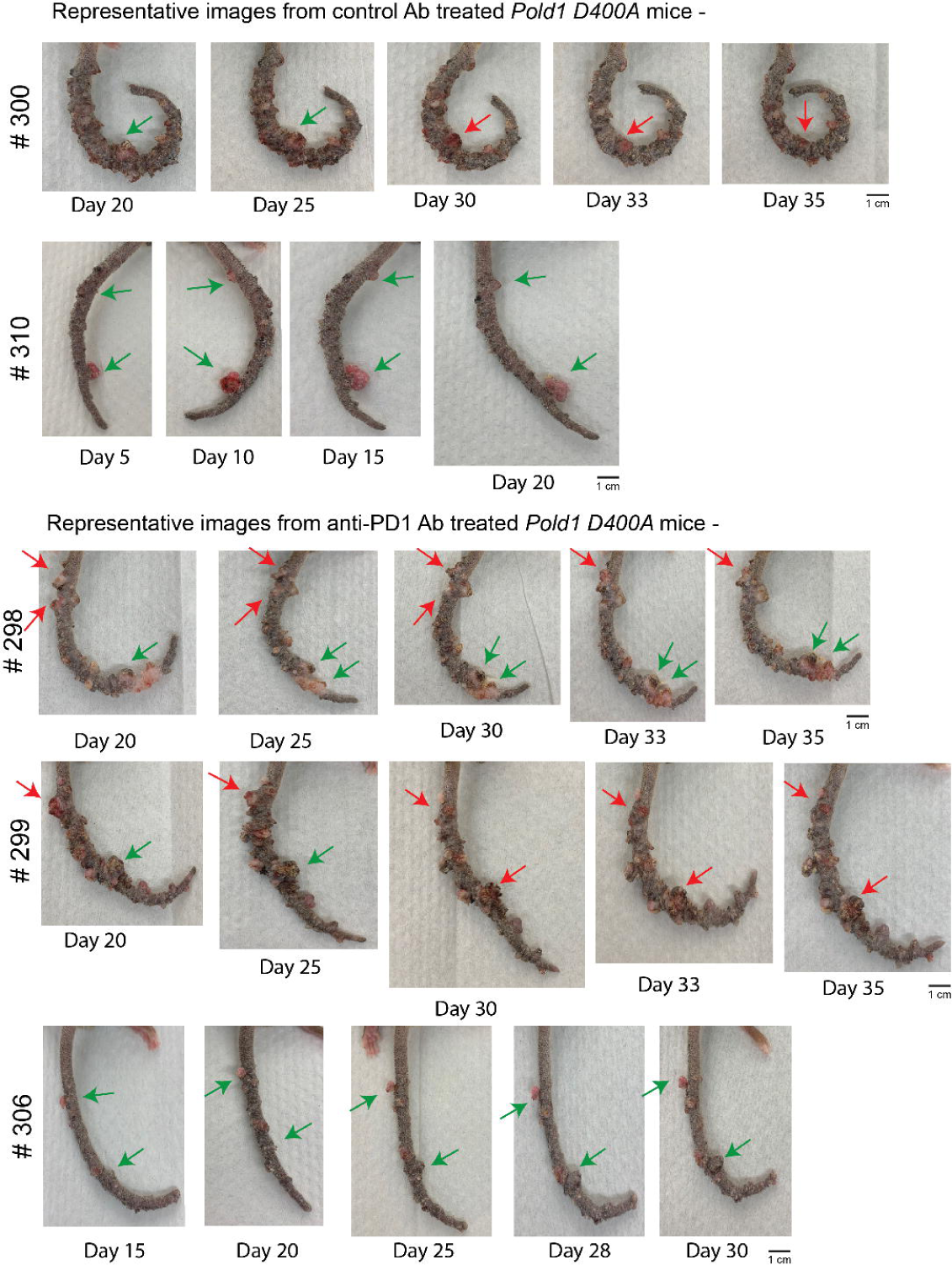
Representative images of tail tumors from *Pold1 D400A* mice were treated with either control IgG or anti-PD1 (RMP-14 clone) antibody every 5 days and tumor size was monitored over their lifetime. The green arrows indicate tumors that remained stable or increased in size. The red arrow indicates tumors that regressed with treatment.

**Supplementary Figure 4.**
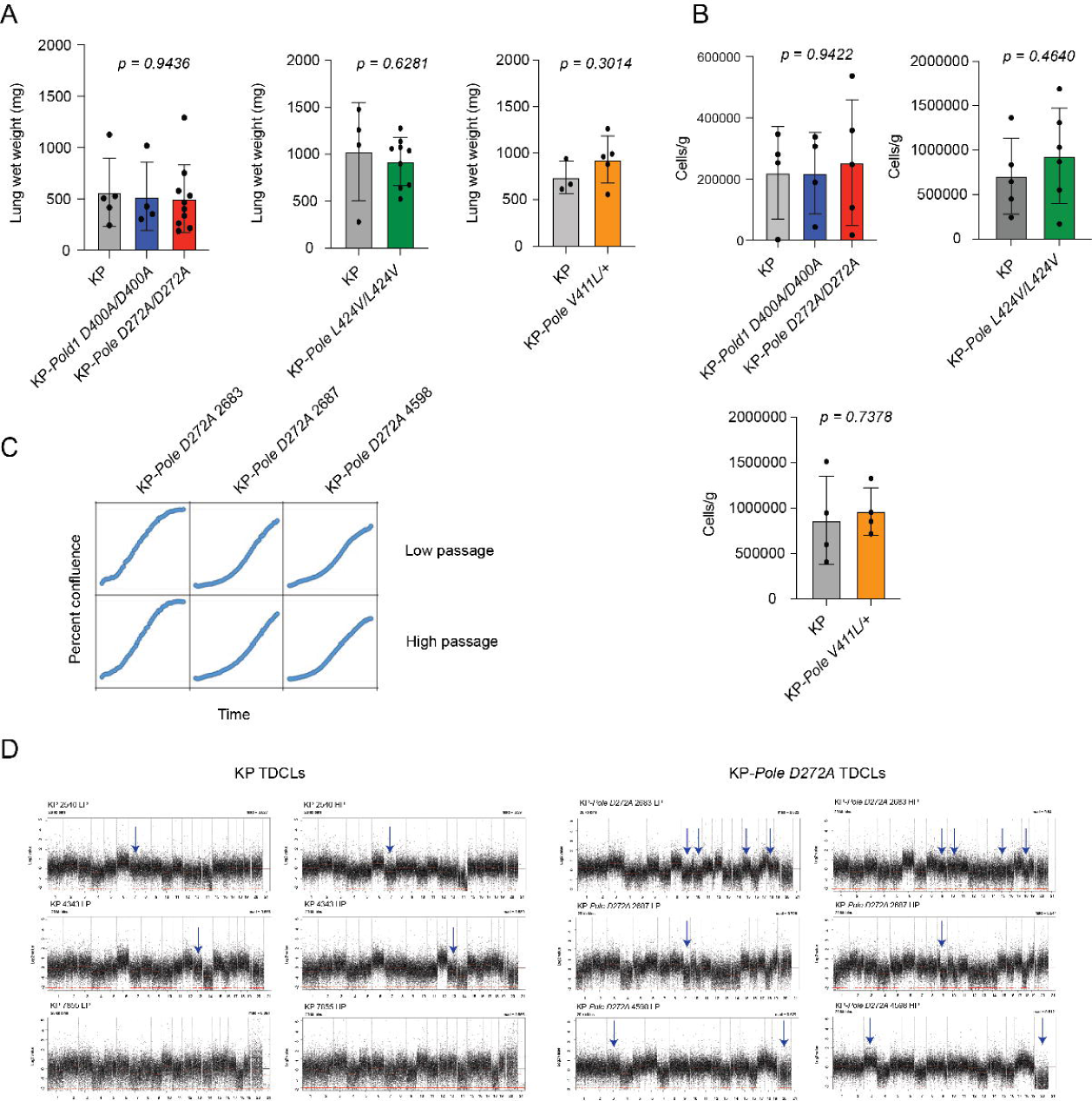
(A) Lung wet weight analysis at 12-week timepoint in KP, KP-*Pole D272A E274A,* KP-*Pold1 D400A*, KP-*PoleL424V* and KP-*PoleV411L/+* mice infected intra-nasally with adeno-cre. *p* values are determined using Student’s unpaired t test. (B) Quantification CD3 positive T cells at 12-week timepoint in KP, KP-*Pole D272A E274A,* KP-*Pold1 D400A*, KP-*PoleL424V* and KP-*PoleV411L/+* mice infected intra-nasally with adeno-cre. *p* values are determined using Student’s unpaired t test. (C) Incucyte growth curve analysis of low and high passage KP-*Pole D272A E274A* cell lines *in vitro.* (D) CNV profiles of KP and KP-*Pole D272A E274A* TDCLs from low and high passage clones. The arrows indicate the chromosomal regions with gross changes.

**Supplementary Figure 5.**
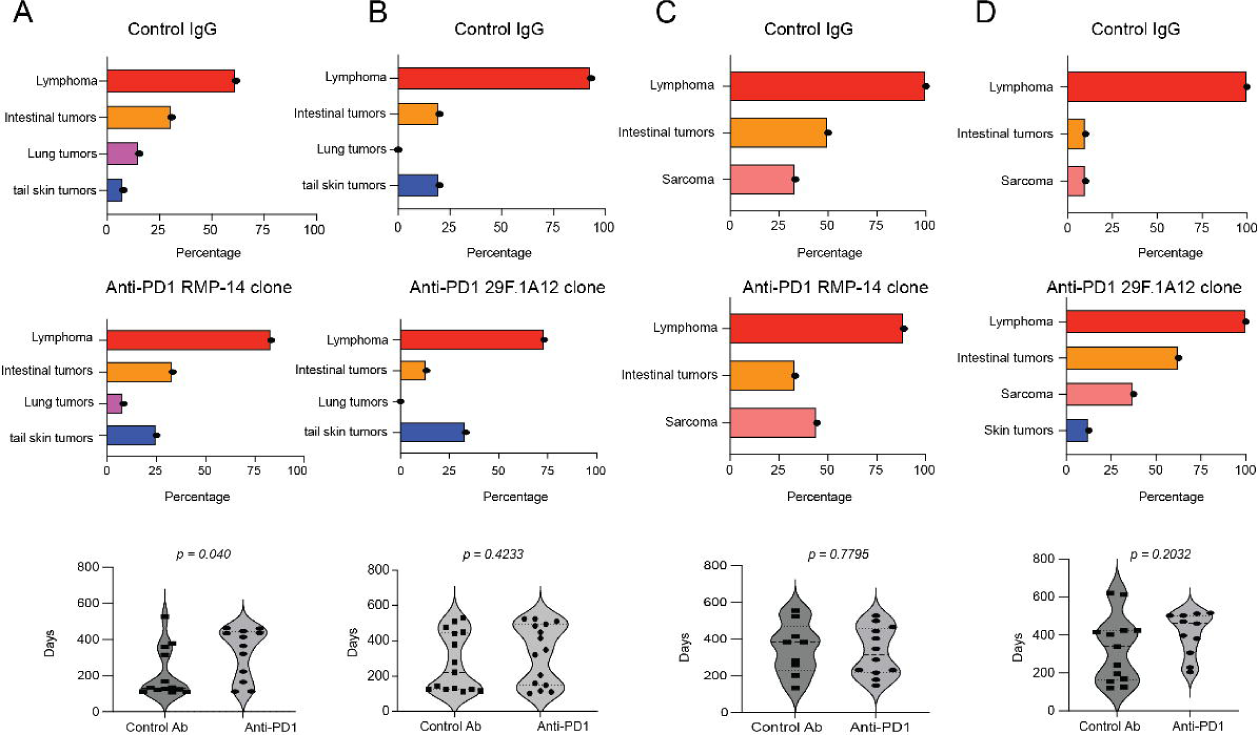
(A) Tumor incidence profile and mean survival of Control IgG and Anti-PD1 antibody (RMP-14 clone) treated *Pold1 D400A* homozygous mutant mice and (B) Control IgG and Anti-PD1 antibody (29F.1A12 clone) treated *Pold1 D400A* homozygous mutant mice. (C) Tumor incidence profile and mean survival of Control IgG and Anti-PD1 antibody (RMP-14 clone) treated *Pole L424V* homozygous mutant mice and (D) Control IgG and Anti-PD1 antibody (29F.1A12 clone) treated *Pole L424V* homozygous mutant mice. *p* values are determined using Student’s unpaired t test.

